# Becoming Biomedical Faculty: An Analysis of Credentials among Successful Academic Career Aspirants

**DOI:** 10.64898/2026.05.20.726576

**Authors:** Cilka M. Hijara, Remi F. Jones, Christine V. Wood, Robin Remich, Anne E. Skelley, Patricia B. Campbell, David P. O’Neill, Richard McGee

## Abstract

Understanding what is requisite for attaining a biomedical faculty career is crucial for guiding trainees preparing for these roles. For nearly two decades, we have collected accounts of biomedical training and career transitions from a large cohort through annual in-depth interviews and tracking of competencies and achievements. This paper elucidates the common and varied credentials of 40 who entered research-intensive faculty careers (RIFCs). Participants completed PhDs and postdocs in a range of research-intensive institutional settings. Developing research independence and a niche were essential to RIFC attainment, and mentors played a crucial role in this development. Counter to common assumptions, high-prestige publications and grants were not in and of themselves necessary for RIFC attainment. Our findings can aid RIFC aspirants and mentors who guide them.

## INTRODUCTION

The career landscape has shifted dramatically over the past several decades, seeing stagnation of tenure-track positions alongside the tripling of postdoctoral positions and PhDs awarded. Although biomedical research training is traditionally focused on preparing trainees for academic careers, only 15% of biomedical PhDs attain any tenure-track academic position in the six years following graduation (1). These shifts have led to increased interest in and speculation about what credentials are required to advance into these competitive roles. This paper examines credentials among those who become faculty.

There are common conceptions about how one becomes a PI, including what publications and awards one must obtain. Yet limited research has studied what is actually required to attain a tenure-track research-intensive faculty career (RIFC). One study that examined requirements in neuroscience concluded that high-prestige products, broadly assumed to play a crucial role in RIFC attainment, are not actually required (2). However, the question of how those without these products demonstrate scientific accomplishment and potential remains open. The findings from another study on training and career transition grants suggest the possibility that a broader range of products might be valued on the academic job market (3).

A small collection of research explores competencies, particularly demonstrations of independence. Some studies categorize different forms of independence, such as performing lab tasks independently and distinguishing one’s research from that of their advisor (4). Another study in the French context links having a distinct research line with increased probability of academic career attainment (5). While such studies establish independence in various forms as a significant part of academic career preparation, a number of questions remain. For example, how distinct must a trainee’s research focus be? How does the process of establishing independence unfold over time? Is independence a requirement for RIFC attainment, or merely an optional competency that betters one’s chances? Overall, research into what is needed to succeed is limited, leaving trainees dependent on approximations of what is essential as they navigate a hypercompetitive academic job market. Thus, it is critical to examine what is requisite for RIFC attainment to guide trainees and mentors as they strive for streamlined, informed, and effective career preparation.

For nearly two decades, our research team has followed a large national sample of biomedical research trainees from undergraduate, postbaccalaureate, or early graduate school years through to the early stages of their careers. Our broader study uses a combination of annual, in-depth interviews and longitudinal tracking of accomplishments and roles. We continue to follow graduates as they navigate the postdoctoral stage and the early stages of their careers.

Using data from 393 interviews with 40 successful RIFC aspirants (see Methods for discussion of sample size), this mixed-methods report examines which accomplishments and competencies enable attainment of RIFCs. Our analysis relies on longitudinal data on research progressions, publications, and grants to examine what is requisite, and what is wrongly assumed to be requisite among those who attain biomedical RIFCs. Our findings provide insight into what is required with the goal of guiding trainees in their efforts toward RIFC attainment. In addition, we hope to inform mentors and leaders in biomedical science so they can more effectively guide trainees toward the requirements of their chosen careers.

By following individuals from the start of training through to attaining a RIFC, we can explore questions such as: How essential are high-profile publications and competitive grants in attaining a faculty position? What types of institutions do successful RIFC aspirants train and work in? What skills and competencies must trainees acquire in order to attain a RIFC? What is wrongly assumed to be requisite? Does timing of accomplishments and competency development matter? What is the role of faculty mentors in the development of essential skills and competencies?

## RESULTS

Becoming an academic research scientist involves the progressive acquisition of knowledge, skills, and competencies. Our data show strong commonalities among the 40 in many aspects of career and scientific development, with nearly all participants engaging in pre-PhD research (n=38); publishing in peer-reviewed journals as first author (n=40); demonstrating research independence at the PhD or postdoc stage (n=37) and finding a research niche (n=37). We also observed variation in institutions attended, publication and grants records, and the timing of acquiring certain achievements and competencies. We include supporting quotations to illustrate participants’ experiences, perspectives, and progressions. All names provided are pseudonyms.

### Characteristics of Colleges and Universities During Training and Faculty Years

We examined whether those who achieve RIFCs train and work in varied or uniform institution types, looking at both Carnegie classifications and amount of NIH funding received. Table 1 provides Carnegie classifications of the undergraduate, PhD, postdoc, and faculty institutions for each participant. Individuals attained RIFCs from undergraduate starting points ranging from baccalaureate colleges to universities with very high research activity. For PhD and postdoctoral training, nearly all participants trained in universities or academic medical centers with very high research activity.

**Table 1.** Carnegie Classification of Participants’ Institutions.

| Carnegie Classification | BS/BA | PhD | Postdoc* | Faculty |
| --- | --- | --- | --- | --- |
| Baccalaureate Colleges - Arts & Sciences | 9 |  |  |  |
| Master's Colleges and Universities (larger) | 3 |  |  | 1 |
| Doctoral/Research Universities | 1 |  |  |  |
| Research Universities (high research activity) | 4 |  |  |  |
| Research Universities (very high research activity) | 20 | 38 | 33 | 34 |
| Doctoral Universities: Higher Research Activity |  |  |  | 1 |
| Special Focus Four-Year: Med Schools & Centers |  | 2 | 2 | 3 |
| Non-degree/non-US/non-Ranked | 3 |  | 5 | 1 |

Cells indicate the number of participants affiliated with each Carnegie institution type at each career or training stage.
\*Four individuals completed multiple postdocs. Counts in this table reflect the most recent postdoc prior to being hired into a RIFC position.

We created a comparative measure of annual NIH funding support which reveals institutional variation beyond Carnegie categories. As described in Methods, we split the range of NIH funding for all institutions represented into eight groups, each consisting of 36 institutions (Table 2). The NIH funding level of participants’ undergraduate institutions varied greatly and included non-research institutions. At the PhD and postdoc stages, almost all trained in the top two groups of research institutions (a range representing 72 institutions). At the faculty level, they redistributed to a wider range of research institutions, according to NIH funding tiers. Thus, this group of successful RIFC aspirants ultimately populated a wide range of research-intensive institutional settings.

**Table 2.**
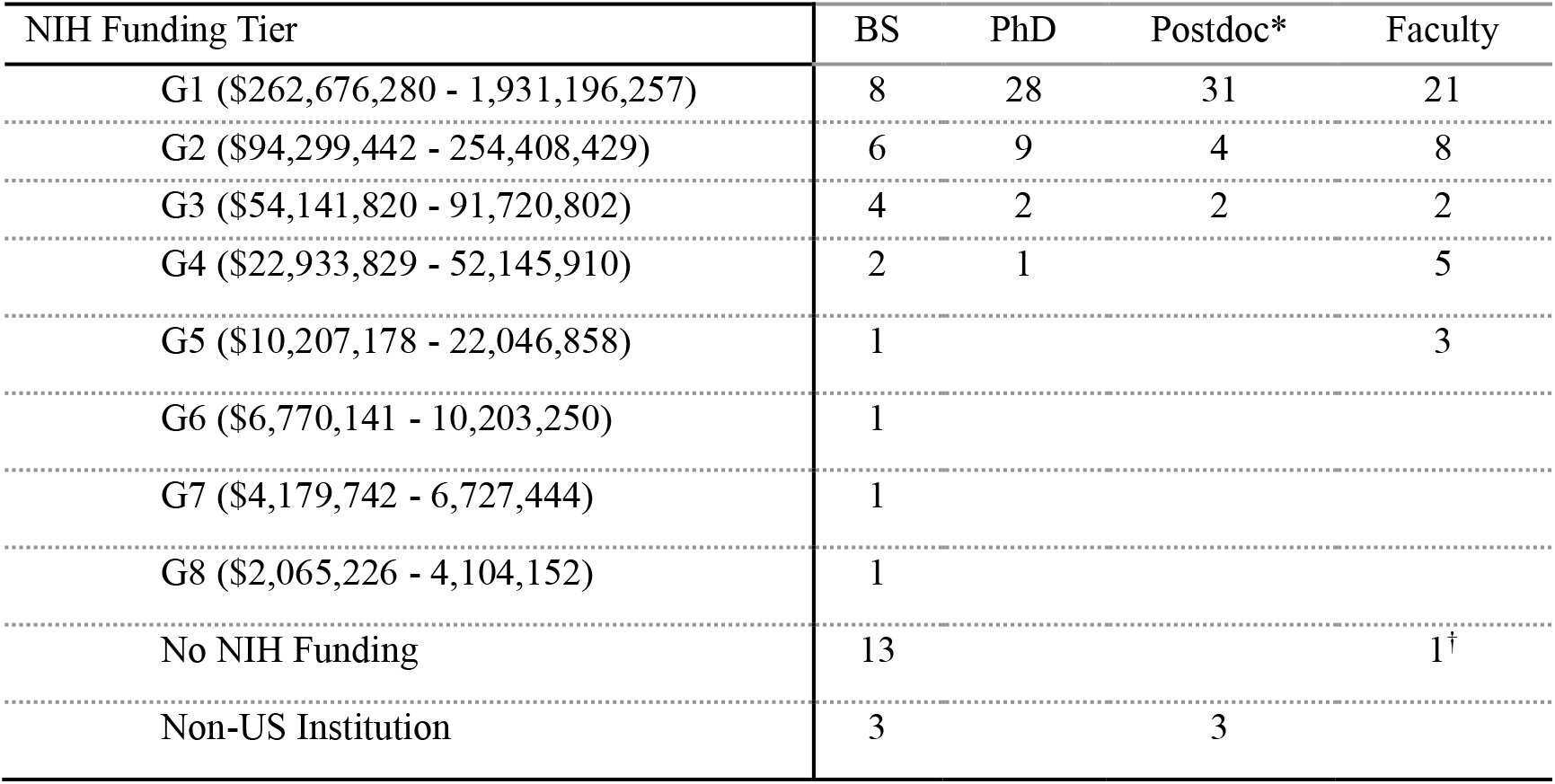
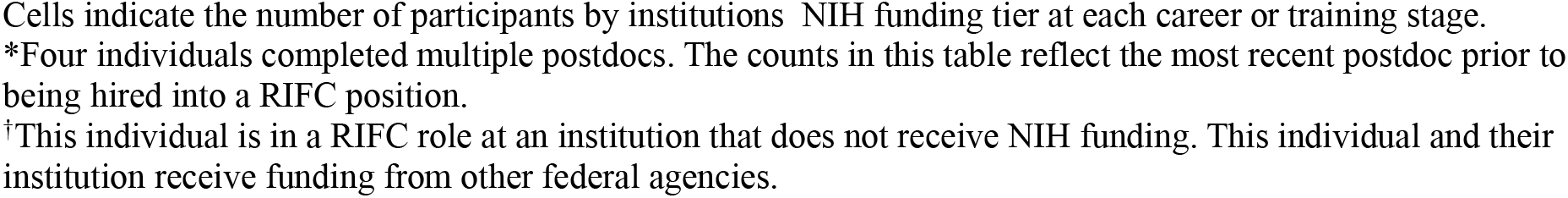
NIH Funding Received by Participants’ Institutions.

| NIH Funding Tier | BS | PhD | Postdoc* | Faculty |
| --- | --- | --- | --- | --- |
| G1 (\$262,676,280 - 1,931,196,257) | 8 | 28 | 31 | 21 |
| G2 (\$94,299,442 - 254,408,429) | 6 | 9 | 4 | 8 |
| G3 (\$54,141,820 - 91,720,802) | 4 | 2 | 2 | 2 |
| G4 (\$22,933,829 - 52,145,910) | 2 | 1 | | 5 |
| G5 (\$10,207,178 - 22,046,858) | 1 | | | 3 |
| G6 (\$6,770,141 - 10,203,250) | 1 | | | |
| G7 (\$4,179,742 - 6,727,444) | 1 | | | |
| G8 (\$2,065,226 - 4,104,152) | 1 | | | |
| No NIH Funding | 13 |  |  | 1<sup>†</sup> |
| Non-US Institution | 3 |  | 3 |  |

Cells indicate the number of participants by institutions' NIH funding tier at each career or training stage.
\*Four individuals completed multiple postdocs. The counts in this table reflect the most recent postdoc prior to being hired into a RIFC position.
†This individual is in a RIFC role at an institution that does not receive NIH funding. This individual and their institution receive funding from other federal agencies.

### Pre-PhD Research Experiences

Nearly all participants (n=38) had formal research experiences before entering PhD programs. All who had research experiences before their PhD programs did so as undergraduate research assistants or in roles between their undergraduate and PhD studies. Post-college research experiences included postbaccalaureate research programs, Master of Science programs, or working in labs in academia.

Those who completed pre-PhD research experiences proactively pursued them to gain essential skills and experiences. Marshall narrated his experience of pursuing research during his undergraduate years:

> I had a really great experience with undergraduate research. I emailed people from a list in the [department], until someone responded, and I started shadowing in that lab the summer after my freshman year, because I wanted to get research experience early. I stuck in that lab all the way through my senior thesis, and I worked there the summer after I graduated, until a couple weeks before coming to [graduate school].

Two participants did not have research experience before starting their PhDs. One decided to pursue a PhD after reading neuroscience literature several years after graduating from college. Another focused on a fieldwork-based scientific discipline for undergraduate study and a first PhD program but did not gain lab research experience until transferring into a second PhD program in a biomedical discipline. In both cases, pre-PhD research experiences were not decisive in determining interest or success in graduate studies. Despite these outliers, early research experiences are key to matriculation into a PhD program and represent an early credential toward RIFC pursuit.

### Publications and Competitive Grants

Scientists seeking RIFCs can make their scientific accomplishment legible on the job market through first-author publications and competitive grants and fellowships. Assumptions about which products are required for RIFCs drive trainees’ publication and grant efforts. Among the most highly cited journals in the biomedical sciences are *Cell, Nature*, and *Science* (*CNS*). *CNS* publications are often regarded as markers of prestige and determinants of career advancement. In our sample, some participants believed that publishing in “the big three” journals was imperative, or even sufficient, for RIFC attainment. Paul said:

…a lot of things will be answered within a postdoc because it’s hard to say that you want a faculty position now. I do want to have a faculty position, but if you don’t get a *Nature, Science*, or *Cell* paper in your postdoc, then those opportunities are not readily available to you anymore.

Similarly, Janelle described her first author “big three” publication as a “golden ticket” to a faculty career.

In reality, there was substantial variation in the timing of publications and the impact of journals in which participants published. While all participants had first-author research publications during their PhDs and/or postdocs, only about a quarter (n=11) had first-author CNS publications.

Publications in journals considered high-impact within a field can also carry value on the job market. We expanded publication records to consider high-impact, field-specific journals (HI-Field) using CiteScore, which indexes by subfield across a four-year window (6; see Methods for description of publication record analysis). First-author, original research publications during PhD and postdoc stages were categorized as HI-Field if they were rated as top 5% in a biomedical subfield on CiteScore.

Thirty participants had HI-Field publications. Ten of these 30 also had *CNS* publications. *CNS* first authorship was achieved most often during postdocs, while the timing of HI-Field first author publications was more evenly split. Nearly a quarter (n=9) attained this career with neither *CNS* nor HI-Field publications. These findings reflect substantial variation in the publication achievements and the timing of publications for those who attain RIFCs.

We also examined participants’ achievement of competitive predoctoral and postdoctoral training fellowships (e.g., F31 and F32) and postdoc-to-faculty career development/transition awards from NIH, such as the K99/R00, and private organizations, such as the Burroughs Wellcome Fund. These awards provide substantial funds for advancing scientific training and starting a RIFC. They are often considered critical for achieving biomedical RIFCs. Nineteen participants did not have a career transition award. Of those 19, six had a competitive training fellowship, and 13 had neither type of award.

PIs and other mentors played an important role in the pursuit of these achievements. In particular, some participants proactively sought guidance in the grant-seeking and writing process. To compensate for her PIs’ limitations and add to her skillset, Melinda formed a relationship with another mentor who had strengths in grantsmanship.

I’m working with a new mentor, supplementary to my current mentor, to help me more with the ins and outs of writing and grantsmanship. It was an eye-opening experience, and I would say it’s a highlight seeing the cell biology community wanting to help and foster young investigators… the mentor that I picked is very much invested in his trainees and it was really inspiring.

Similarly, others recognized the potential shortcomings of their PIs for resource limitations and lack of experience but proactively sought other mentors to receive customized support in areas where their PIs could not be helpful. Branden set up a “secondary” relationship to assist with grant writing, even though he had a positive relationship with his primary PI.

My primary mentor is junior faculty. And then I have a secondary appointment with a much more senior person… I think he’ll end up sponsoring me for grants to improve my likelihood of getting them.

By building a broader network to develop specific skillsets or, in some cases, to compensate for a lack of mentorship in a particular area, participants strengthened their abilities and chances of acquiring competitive products.

Given the surprising number of participants who did not have major publications (n=9) or training/faculty transition awards (n=13), we examined publications and grants in tandem to determine which types of products appeared to be sufficient for RIFC attainment, either alone or when paired with another type of product (see Table 3). Twenty-one participants had a combination of a major publication (CNS and/or HI-Field) and a training and/or transition award, likely to be perceived as a high level of accomplishment. Only 1 of these 21 participants had all major publication and grant types (Table 3,green), reflecting the rarity of this ideal scenario. Notably, only 1 participant attained a RIFC with a CNS publication alone, countering misconceptions that a CNS publication alone is a reliable pathway to securing a faculty role.

**Table 3.** Combinations of Publications and Grants/Fellowships. The table displays participant counts for various combinations of publication and grant/fellowship types attained prior to the faculty stage.

|  | CNS+HI-<br>Field | CNS only | HI-Field only | Neither |
| --- | --- | --- | --- | --- |
| Trans.+Training | 1 | 0 | 3 | 3 |
| Transition only | 5 | 0 | 6 | 3 |
| Training only | 3 | 0 | 3 | 0 |
| Neither | 1 | 1 | 8 | 3 |

Nearly half of participants (n=19, Table 3,dashed red box) attained RIFCs with only a training or transition award, only a major publication, or with neither. Of these 19, nearly half (n=8) had only a HI-Field publication, suggesting that HI-Field publications alone can enable RIFC attainment. Over a quarter (n=6) had faculty transition awards but no major first-author publications; none from this group had training fellowships alone. Like HI-Field publications, transition awards appear to have independent influence on RIFC attainment. Only 3 participants had neither major publications nor training/faculty transition awards (Table 3,blue). In these cases, individuals were in unique institutional settings, or research subfields where other products (e.g., patents, unique methodologies) are highly valued. Overall, our data suggests that individuals can attain RIFCs with different combinations of major publications and grants, and that there is no single type of publication or grant that is required.

### Developing Independence

For each participant, we observed the continuities and change in research independence across PhD and postdoctoral training. We define research independence as a combination of skill, opportunity, and desire to design and conduct original projects and experiments and to achieve autonomy over time and work in the lab. Thirty-seven of 40 individuals demonstrated high independence during at least one of the PhD or postdoc stages, reflecting an effort to achieve independence alongside the opportunity to enact it. More than half (n=23) demonstrated high research independence during graduate school, with all but one increasing or maintaining independence throughout their postdocs (Figure 2).

**Fig. 1.**
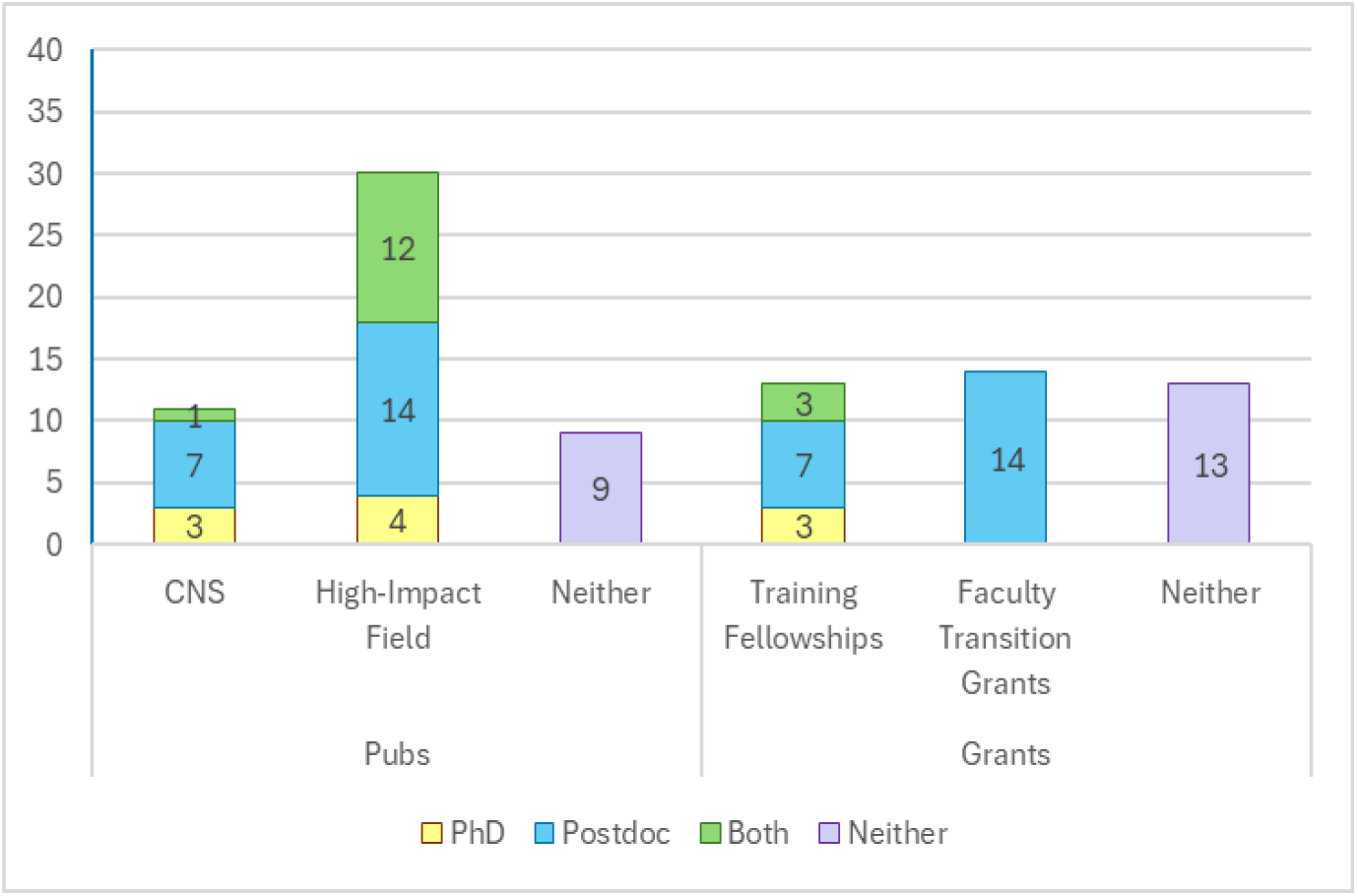
Timing of Publications and Grants/Fellowships. Bars display counts for publication and grant/fellowship categories. Color coding represents training stage during which publications and grants/fellowships were attained.

**Fig. 2.**
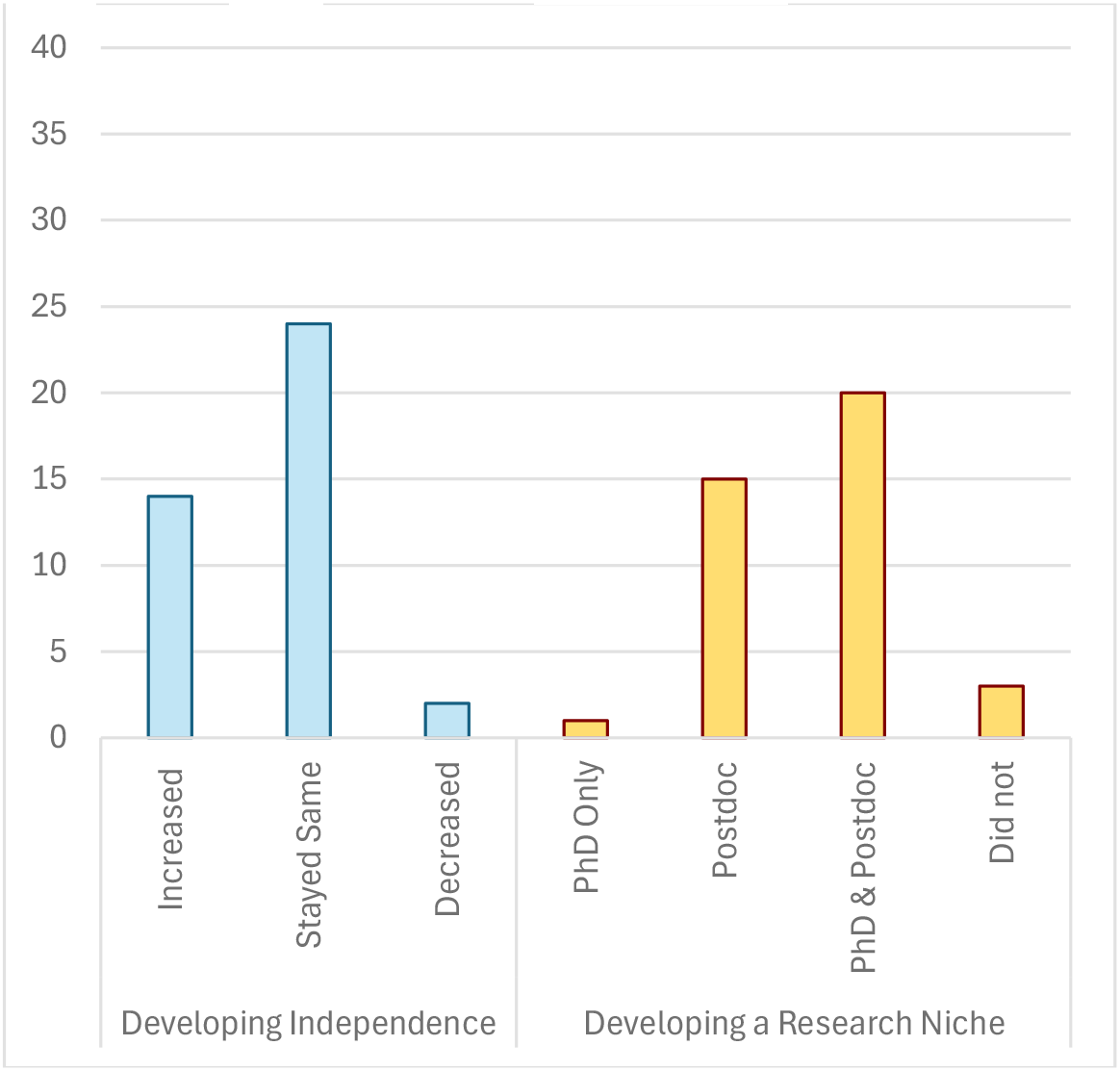
Competencies Developed During PhD and Postdoctoral Training. The figure displays both the number of participants fitting three patterns of independence over PhD and Postdoctoral training as well as when participants developed their research niche.

Narratives of growing independence often focused on achievements and gradual shifts such as successfully navigating a PI’s limited availability, growing to rely less on PI guidance, gaining decision-making power over experiments and projects, designing new projects and receiving permission to pursue them, and strategizing ways to advance independence. While the typical trajectory among this sample was to increase or maintain a high level of independence, some participants experienced instances of limited independence. Participants report that developing independence is often influenced by a PI’s approach to training. PIs can hinder or nurture independence through their interactions with mentees and through the research environments they foster. Below, we display how participants contextualized instances of high and low independence.

Christopher’s two PhD PIs were available but allowed him to work independently from an early stage.

> I feel very independent. I really don’t talk to either of my PIs very much, except when errors come about [or] I have pressing questions… I choose the directions to go in. I choose the systems and the things that I want to look at.

Dustin took his postdoc PI’s busyness as an opportunity to increase his independence, which he believed would benefit his career.

> …my advisor is a director of a huge institute, so he doesn’t have much time to direct individual people. That can be a problem for a lot of people if they can’t navigate that amount of independence, but for me it was good to struggle through that… So, I feel a sense of ownership of where I’m going and what my identity as a scientist will be in the future as well as now.

During her PhD, Nikki felt micromanaged by her PI. During her postdoc, her independence grew as her PI encouraged work on her own ideas and experiments.

> There was this one meeting that I had with my [current] boss, and I pitched this experiment to him before I went and ordered any reagents or anything. And he just looks at me, and he’s like, “You’re a postdoc, go do it.” I was like, “Oh yeah, I guess I am.” There’s a lot of confidence-building that comes with that… having to own all my decisions.

Among those demonstrating low research independence at any time, participants described their PIs assigning their projects or controlling their work. Jeffrey described his postdoc PI as overly demanding.

> [My PI] would tell me I can do this more independent project, and then I’d start working on it, and then she’d tell me I can’t. She would take on all these projects and all these responsibilities and just start dumping things on [me].

Not all participants attributed their independence to their PIs. Laurie felt she needed more confidence in bringing a project full circle; she sought opportunities to build confidence and gain independence during her postdoc.

> I need to gain the confidence that I can come up with a project and lead a project and publish it from start to finish, because in my PhD I did have independence, but I wasn’t necessarily from start to finish coming up with the idea of the project, so I’m really looking and hoping to gain that kind of confidence.

Developing and demonstrating research independence is a key competency toward becoming and performing the role of a PI. Nearly all participants described consistently high or increasing independence over the course of training. However, some participants experienced and overcame instances of limited independence. Participants articulated an interplay between their desire for independence and their PIs, who created structures and environments in which independence thrived or was stymied.

### Developing a Research Niche

Developing a niche—a specialized and well-defined research focus—is an important aspect of scientific development, distinct from developing research independence. Through a niche, scientists define a distinct line of research and contribute original knowledge to their fields, gaining legitimacy among their peers. Many participants did not develop a niche during graduate school, because their projects were so closely related to their PIs’ work, even if they were operating independently. More than one third of participants (n=15) first developed a niche during postdocs when articulating a unique and portable line of research becomes more important for representing and distinguishing oneself on the academic job market (Figure 2). The timing of niche development among this group suggests that, for RIFC aspirants, the postdoc serves as a unique and significant stage in scientific progression.

Those who developed a niche during the PhD (n=21) tended to follow that line of research and refine it further during the postdoc. During his PhD, Marshall worked under the direction of two PIs to develop a niche.

> My research kind of bridges the two labs. [One is] more of a molecular lab… we make [pathogens] that have interesting properties, and then we characterize them. It’s very much focused on the [pathogen]. The other lab is… more focused on the immune response. My project is kind of in the middle, taking these interesting [pathogens] and [developing treatments].

He sustained this interest throughout his PhD, eventually narrowing to specific pathogens and continuing to study them through his postdoc.

Developing lines of research to take to their future labs was a key element of niche development. This process depended on trainees’ willingness to take risks, and their PIs’ willingness to let them try new lines of inquiry. For Jessica, this meant developing a novel application of imaging technology to a different disease model for her future lab.

> A big part of my postdoctoral journey was developing a way to connect the images that we were generating with the gold standard of the field…I asked [my PI], “Can I try this for [disease]?” And he said, “You’re not going to see anything.” And I said “Yeah, but can I try it?” And he was kind enough to let me try it, and we started seeing all sorts of crazy things, and I just got hooked on it.

For some, niche development was paired with an awareness that not all niches are equally fundable. Bruce worked primarily on developing a “marketable” postdoc project that would help him find a faculty position, rather than simply pursuing specialized expertise. Bruce said:

> I had a friend who is looking for a faculty position now, and he said, “Pick the project that would be more marketable in three or four years, like when you start to look for jobs, and really make sure that you’ll be able to take [it] with you, as your project.”

Niche development and articulation are active and strategic processes. Participants reflected on how their niche would fit into career plans, both in terms of becoming a line of research for a RIFC and appealing to potential employers and colleagues.

Niches typically branched off from PIs’ research foci, representing iterations rather than dramatically different lines of research. We followed iterations of niche development for individual participants over time and observed the following sequence: building expertise in a PI’s research focus, exploring a new branch or line of inquiry, and negotiating ownership of this branch with the PI. For Ariana, developing a niche entailed finding an area of research she was passionate about and gradually focusing on more specific questions she hoped would produce novel findings. She also narrated hurdles with her PI and lab dynamics that slowed her process of claiming a niche. Early in her postdoc, she said:

> I’m willing to work hard and apply for funding opportunities. The problem is that [it] requires having a project. At the point where I’m at, I don’t have the freedom to choose my own project and to go for it. I kind of have to wait for something that is given to me, and I’ll work with that, and if something comes from that, I could take [it with me] as a project. And so, at this point, I’m probably not very competitive because I don’t have an independent project.

Over the next several years, Ariana published novel findings that made her feel closer to having her own project. However, she still worried about differentiating her work from her PI’s and was unsure of which research lines were hers to take. When Ariana shared these concerns, the interviewer pointed out that she had made a novel discovery of her own and questioned why she would be worried about claiming this as her niche. Ariana explained:

> It’s done in his lab, right, and so I think it’s a conversation. But even if you have a conversation of, “Well I’m going to work on it,” it doesn’t mean that somebody else in his lab [won’t] come and work on it, because it’s happened before. So yeah, those are just things that I didn’t consider but that I need to think [about] going forward.

Later in her postdoc, Ariana received a K99/R00 award, which empowered her and served as a catalyst toward refining a line of research. She viewed the K99/R00 period as an opportunity to “potentially [create] another field of study” that she would carry with her into the faculty position. Ariana’s experience trying to differentiate her work from her PI’s demonstrates the complexity of finding a niche.

All but three participants developed a research niche by the end of postdoctoral training. These three instead developed specialized technical skills or methodologies that could be applied to different fields, in effect demonstrating technical or methodological niches. These exceptions show the high valuation of technical innovation in related subfields such as biomedical engineering. Overall, these results demonstrate that those who became PIs uniformly demonstrated scientific potential through the development of a distinct niche.

## DISCUSSION

Becoming a PI involves years of scientific training, professional development, and demonstrating accomplishments. The results in this article shed light on what is requisite to achieve this career, including competencies such as research independence and achievements such as first-author publications. Our findings also identify credentials that tolerate variation, such as publication types and institutions attended. Additionally, our findings illustrate the important role that mentorship plays in acquiring these requisite accomplishments and competencies.

Demonstration of scientific accomplishment is essential for career attainment but can take different forms. One question in the biomedical field is whether publishing papers in the highest tier journals (*CNS)* and/or obtaining faculty transition grants during the postdoc are critical to attaining a RIFC. Previous literature has examined rates of K99/R00 awards and *CNS* publications in neuroscience, finding that neither are necessary for obtaining tenure track positions (2). In our sample, only 11 participants had *CNS* publications prior to starting RIFCs. We examined a broader set of publications which included both *CNS* and HI-Field (high-impact field-specific) publications and found that 31 participants had one or both. Additionally, we expanded the view beyond the K99/R00 to include private foundation grants and competitive training fellowships. We found that faculty transition grants are common (n=21 of 40) among those who obtain RIFCs, and that most participants received transition and/or training awards (n=27 of 40).

Fourteen participants had neither a transition grant nor a *CNS* publication, revealing that certain accomplishments are not absolute requirements. They found other avenues to RIFCs, most commonly by publishing in disciplinary journals that had high impact factors in their fields. In light of this finding, RIFC aspirants and PIs might focus their publishing strategies on an array of journals that includes high-impact, field-specific outlets. A small number of participants had training fellowships paired with high-impact publications. However, unlike HI-Field publications, training fellowships do not appear to influence RIFC attainment alone. Dissemination of information to trainees and mentors about which products are crucial, and which are valuable but optional, would aid in streamlining career preparation.

Our findings also identify requisite competencies. Once trainees enter graduate school, they continue a process of scientific and professional development that involves cultivating skills and competencies that are essential for successfully performing the role of a PI. These include developing both research independence and a research niche. By examining patterns of independence and niche development among a population of successful RIFC aspirants, our work establishes that these competencies are requisite, rather than optional-but-advantageous skills.

For more than half of participants, independence was distinct from and acquired before identifying a niche, with some arriving at a niche late in their postdocs. That this process occurs or continues during the postdoc years suggests that the postdoc represents a distinct and crucial stage of preparation for those pursuing RIFCs. Additionally, developing a niche was an iterative process that typically involved emulating and building on a PI’s research focus, as opposed to identifying a radical new direction.

Previous work has pointed to research independence as a key factor motivating PhD students early in their trajectories to choose to pursue and continue in research (7). The vast majority of participants either maintained a high level of independence or increased their level of independence over the course of training. Participants described independence as the result of desire, skill, and opportunity. PIs played a crucial role in providing opportunity, complementing literature that emphasizes mentors’ important role in cultivating independence among mentees (8). Formalizing and clarifying the related paths towards both independence and a niche would make mentoring practices more effective and better prepare trainees who seek RIFCs.

Our focus on a subpopulation that attained RIFCs enables a detailed within-group comparison of credentials acquired during training but excludes comparison with those who sought this career and did not attain it. In the broader NLSYLS population, there are 52 participants who had RIFC intentions at the time of graduation but did not attain this career. An in-progress comparative study of these individuals will provide further clarity on the contours of achievements, competencies, and skillsets that allow some individuals to attain RIFCs but not others.

Finally, while this paper explores what is requisite for attaining a RIFC, there is a parallel need for understanding what is requisite for success once one attains a RIFC. With individuals in our population followed through the first years of their faculty careers, we will explore what shapes success in processes such as grant-seeking and tenure for early-career faculty.

## MATERIALS AND METHODS

### Description of the Population and Data Sources

Study participants came from two NIH-funded longitudinal studies investigating career preparation, decision-making, and differentiation among a national population of biomedical graduate students. Seventeen of 40 subsample participants were enrolled in the National Longitudinal Study of Young Life Scientists (NLSYLS). The NLSYLS is an interview-based study of career decision-making among biomedical research trainees from the beginning of the PhD through the early career stage. Two hundred sixty participants were recruited from colleges and universities across the US and Puerto Rico in multiple waves from 2008 to 2011, either as beginning PhD students or as late-stage undergraduates/postbaccalaureate students with intentions to enter graduate school. Using tracking and annual interviews, the NLSYLS followed trainees until they differentiated into non-academic careers, opted to discontinue participation, or left their PhD programs.

Twenty-three participants in this subsample are from a second interview study, The Academy for Future Science Faculty (“the Academy”). Starting in 2011, The Academy tested the effectiveness of a theory-driven career coaching intervention on persistence toward RIFCs. The Academy recruited cohorts of PhD students at US universities; half at the start of PhD programs, and half nearing graduation from their PhD programs. Participants at each stage were randomized into control and experimental groups.

Experimental groups received virtual and in-person coaching from senior biomedical faculty with extensive mentorship experience for up to three years. Thirteen participants in this study were in control groups, and 10 were in experimental groups. All participants, control included, completed annual interviews focused on both their experiences in the Academy and their experiences in graduate school and at subsequent stages of training. This report draws only on interview and tracking data; an evaluation of the Academy intervention has been reported previously (9).

An R35 MIRA award allowed the team to merge the Academy interview data into the NLSYLS project. The data from both studies are treated equivalently, because data collection approaches were nearly identical. Questionnaires are provided in a previous publication (10). Additionally, we used identical procedures and publicly available data sources to track career outcomes (faculty/staff pages, Linkedin), publications (PubMed, Google Scholar), grants (NIH RePORTER), and categorize training and faculty institutions (NIH RePORT).

The population of 40 represents most of those across the two studies who obtained RIFCs. A total of 54 participants across both studies obtained RIFCs but 14 are not included in this report because sufficient interview data were not available for a thorough analysis. All participants are identified using pseudonyms.

Table 4 provides the sociodemographic characteristics of the group. The group contains an equal number of men and women, although the total population in the NLSYLS consists of 61% women. While Black/African American and Hispanic individuals each make up 12.5% of the 40, a proportion higher than new faculty hires in the biomedical sciences (8), our overall study population of doctoral recipients is 17% Black/African American and 18% Hispanic, much higher than the respective 3% and 9% reported by the NSF (2). Individuals in this group came from varied socioeconomic settings, with 13 of 37 who provided these data growing up with family incomes below the mean household income. In contrast, 15 of 37 fell into the two highest income categories. Twenty-five percent had parents with a high school education or less, and 25% with a bachelor’s. Finally, four were first-generation immigrants and five were children of immigrants. Thus, our sample includes successful RIFC aspirants from varied backgrounds.

**Table 4.**
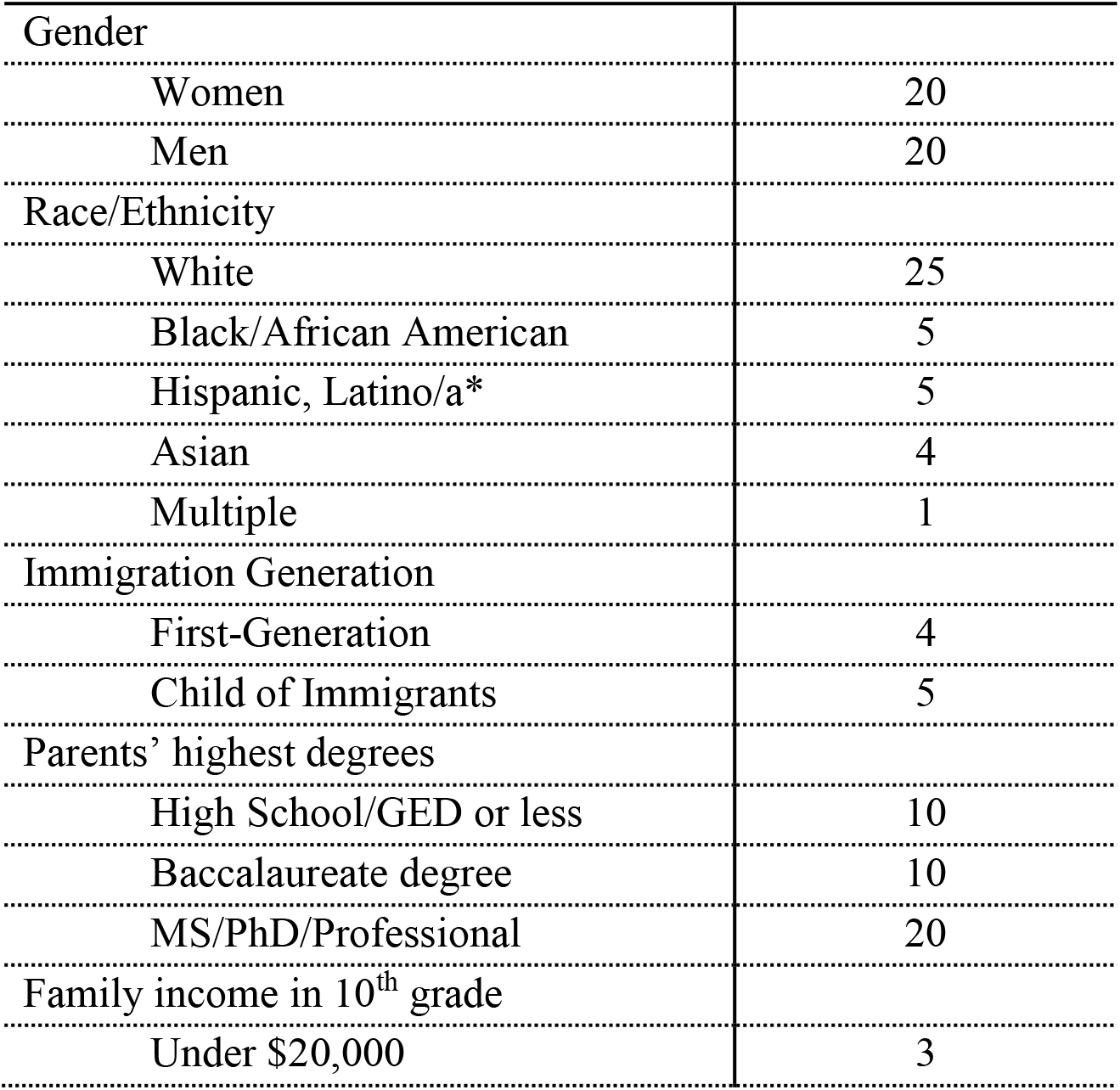
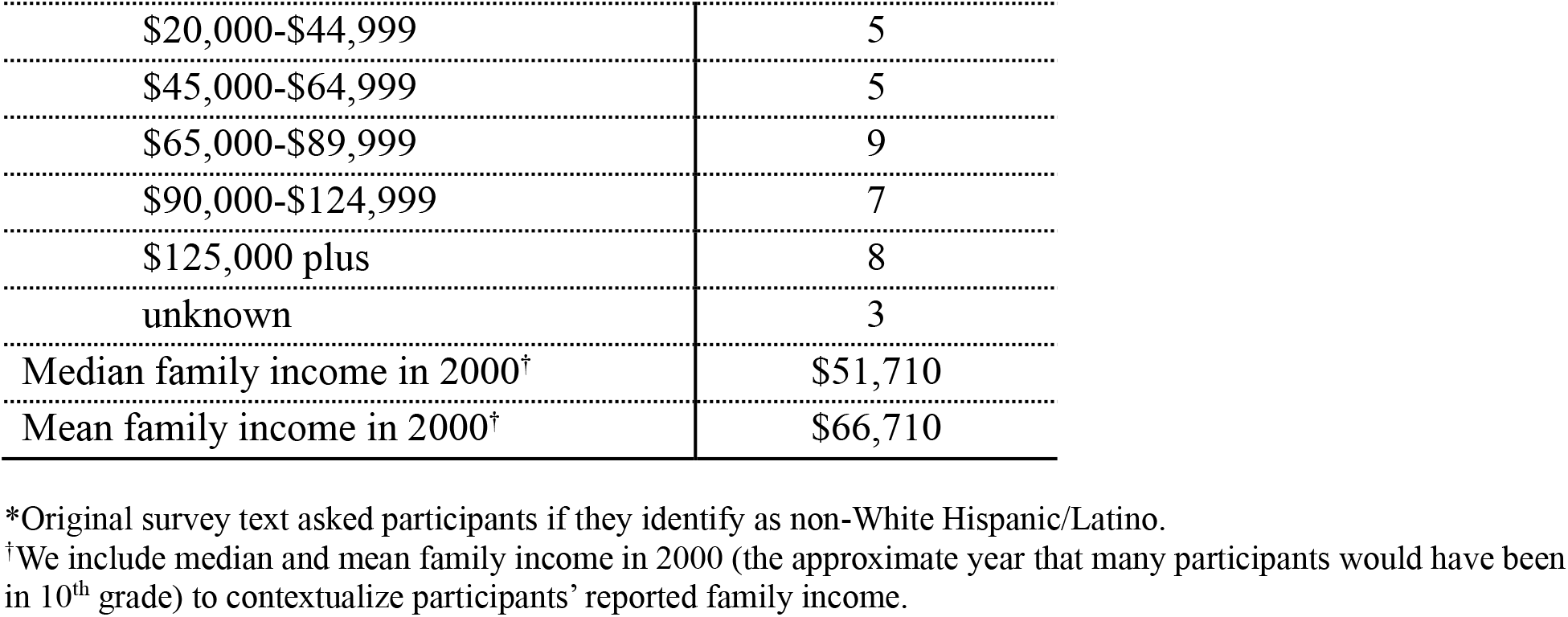
Participant Sociodemographic Characteristics.

### Data Collection and Analysis

Data collection and analysis focused on a combination of annual qualitative interviews and tracking of publications, grants, and institutional affiliations. Qualitative interviews took place in person or by telephone or videoconferencing. Interview guides contained questions about academic and research experiences prior to and during graduate school; mentors and PIs; and career thinking and preparation. Unlike many studies of career preparation and decision-making among biomedical scientists which use cross-sectional approaches, our data reflects participants’ responses in real time elicited at regular intervals

Our sample size (40 participants, 393 interviews) and analysis strategy achieved saturation, a metric for judging completeness and adequacy of sample size and analysis in qualitative research. Saturation is defined as the point at which no additional themes or patterns are found, and researchers are empirically confident that conclusions are thoroughly developed (13-14). Saturation is typically achieved early in analysis and at small sample sizes (e.g., 9-12 interviews; see 14-15). We were able to identify patterns, their meanings, and their importance after developing and discussing summaries of about 10 participants. However, we continued beyond the point where we reached saturation to ensure that our conclusions were based on the widest possible range of participants, and to examine where variations fell along demographic lines.

The research team coded interview transcripts using NVivo qualitative software as previously described in depth (9). Team members used coding reports and close readings of full transcripts to create detailed longitudinal summaries for each participant focused on research progression, breakthroughs, and challenges. Summaries included a detailed history of each participant’s experiences before enrolling in a PhD program through their postdoctoral work and eventual job selection, along with supporting quotes. Each summary was written by an individual team member and discussed by the team to identify areas that needed expansion, clarification, or further expansion. This approach promoted consensus across multiple perspectives. From these summaries, the team performed secondary analysis to identify commonalities and variation across the sample.

To examine patterns in credentials prior to attaining a RIFC, members of the team (CMH, RFJ, RM, and RR) created a spreadsheet with categories capturing competencies and skills from an initial set of summaries. The initial set of categories included competencies such as project management and supervising, mentoring and teaching, collaboration, grant writing, developing research independence, and developing a research niche. Ultimately, we focused on developing research independence and developing a research niche, as the paths toward these competencies were individualized, not an inevitable result of the training structure, and were attached to clear demonstrations of scientific accomplishment in the job-seeking context. The full team explored these competencies and populated the participant fields with fixed-response categories (e.g., binary responses such as “yes”/”no”). Additionally, we added short text summaries to accompany fixed responses for each category to give meaning and context. We assessed independence and niche development separately at both the PhD and postdoctoral training stages so we could track progressions and changes over time. This strategy allowed us to capture temporal information such as increases, declines, and stability in independence.

A separate analysis focused on grant and publication records. A first round of grant records was compiled using NIH RePORTER (16). Specifically, we recorded when participants received competitive training fellowships (F31, F32) and faculty transition grants (K99/R00), based on common conceptions of the advantage of having these awards in RIFC-seeking. A second round of grant records focused on faculty transition grants funded by private institutions, which are comparable to NIH K99/R00s. These grants include Burroughs Wellcome Fund and Hanna H. Gray Fellows Program and were self-reported by participants when they were asked about the past year’s accomplishments at the start of each interview. After compiling grant records, we distilled them into binary categories (yes/no) that captured if they had received each grant type, and at what stage of training (PhD or postdoc).

We compiled publication records using PubMed (17). Our analysis included first-author, original research publications received prior to attaining a RIFC. We removed all review articles, patents, and abstracts from each participant’s list of publications, along with all articles published after the start of faculty positions. Additionally, we checked each publication where a participant was listed in the first 5 author placements for co-first authorship.

Once we had compiled a list of first-author, original research publications received during training, we searched publication lists for two categories related to the prestige and impact of the publishing journal: *CNS* (*Cell, Nature*, or *Science*) and HI-Field (high-impact field-specific outlet). HI-Field publications were operationalized as any first-author, original research article that appeared in a journal listed in the top 5% in a biomedical subfield on the Scopus CiteScore index. Each journal indexing system offers advantages and disadvantages. We favored CiteScore based on its inclusion of a broad range of outlets and its use of a relatively wide 4-year citation window. As with grants, we distilled our final set of records into binary (yes/no) categories reflecting if participants had received each publication type during each stage of training.

Finally, we analyzed the institutions where each participant completed PhDs and postdocs or held faculty positions to determine whether those who achieve RIFCs train and work in varied or uniform institution types. First, we sorted institutions using the Carnegie classification system. The Carnegie system displayed the variety of undergraduate institutions but collapsed PhD and postdoc stage institutions largely into one category. Therefore, we created a comparative measure of annual NIH funding support to capture variation among research-intensive institutions. To create this comparative measure, we split the range of NIH funding (publicly available on the NIH RePORT website) for all institutions represented into eight groups, each consisting of 36 institutions (Table 2), excluding institutions with less than $2 million in NIH funding (no participant had an affiliation with one of these institutions). Initially, we split institutions into 10 groups but found that a number of highly regarded and highly active institutions were arbitrarily sorted into the second funding tier due to outliers at the upper end of the NIH funding range.

The goal of analysis of competencies (independence and niche), accomplishments (publications and grants), and institution type and funding tier was not only to find common credentials, but also to identify those which tolerated variation, especially in cases where patterns ran counter to common assumptions. An initial step was to compile counts of fixed choice responses and identify themes with consistency and variation. Qualitative analysis involved reviewing short response entries and summaries to provide further insight into how participants perceived and achieved key competencies and accomplishments. The results describe the credentials (competencies and accomplishments) that shape attainment of biomedical RIFCs.

## Acknowledgements

We would like to acknowledge and thank the many individuals who have worked with our team since the beginning of the longitudinal studies in 2008. First, Dr. Jill Keller, who was a key partner in initiating the studies and the qualitative research expert at the beginning. Others who played invaluable roles at different stages of the project to whom we are indebted include Fatimah Bhatti, Bryan Breau, Adriana Brodyn, Lynn Gazley, Toni Gutierrez, Sandy LaBlance, Nicole Langford, Ebony McGee, Elizabeth Morrissey, Michelle Naffziger-Hirsch, Letitia Onyango, Jennifer Richardson, Bhoomi Thakore, Simon Williams, Veronica Womack. We would also like to thank Dr. Jeff Engler for his valuable suggestions on an early draft of this manuscript.

## Funding

This work was funded by grants to R.M. from:

National Institute of General Medical Sciences grant R01 GM085385

National Institute of General Medical Sciences grant DP4 GM096807

National Institute of General Medical Sciences grant R01 GM107701

National Institute of General Medical Sciences grant R35 GM118184

National Institute of Nursing Research grant R01 NR011987

## Author Contributions

Conceptualization: R.F.J., C.M.H., C.V.W., P.B.C., R.M.

Methodology: R.F.J., C.M.H., R.R.

Software: R.F.J., C.M.H.

Validation: R.F.J., C.M.H., C.V.W.

Formal analysis: R.F.J., C.M.H., C.V.W., R.R., D.P.O’N., R.M.

Investigation: R.F.J., C.V.W., R.R., P.B.C., A.E.S., R.M.

Data Curation: R.F.J., C.M.H.

Writing - Original Draft: R.F.J., C.M.H., C.V.W., R.M.

Writing - Review & Editing: R.F.J., C.M.H., C.V.W., R.R., D.P.O’N., P.B.C., A.E.S., R.M.

Visualization: R.F.J., C.M.H.

Supervision: R.M.

Project administration: R.M., R.F.J., C.M.H., C.V.W., A.E.S.

Funding acquisition: R.M., P.B.C.

## Competing Interests

The authors declare that they have no competing interests.

## Notes

### Competing Interest Statement

The authors have declared no competing interest.

